# Metabolic tagging of adipose-derived stem cells for targeted modulation of regenerative potency within mineralized collagen scaffolds

**DOI:** 10.64898/2026.09.25.754411

**Authors:** Jaejung Kim, Dhyanesh Baskaran, Alison C. Nunes, Génesis Ríos Adorno, Hua Wang, Brendan A.C. Harley

## Abstract

Recent efforts to reconstruct critical-sized craniomaxillofacial defects using biomaterials have increasingly sought strategies to accelerate proangiogenic processes immediately after surgical implantation to improve host-graft integration. A class of mineralized collagen scaffolds has recently been identified to promote osteogenesis of bone marrow and adipose-derived mesenchymal stem cells both *in vivo* and *in vitro*, motivating efforts to identify strategies to promote osteogenic-angiogenic coupling in order to support regenerative healing. Here, we report the first-time use of a glycoengineering approach to generate a population of engineered ASCs (eASCs) from human adipose-derived stem cells (hASCs). This effort inserts an azido sugar onto the cell membrane that can be selectively targeted using DBCO-tagged biomolecules to selectively boost eASCs activity. We report the effects of azido sugar labeling conditions (concentration and incubation time) on eASCs labeling efficiency. We show proliferation and pro-regenerative gene expression activity of eASCs in mineralized collagen scaffolds are largely similar to conventional hASCs. And we demonstrate that molecular cargo functionalized with dibenzocyclooctyne (DBCO) can be conjugated to azido groups on eASCs via click chemistry. Together, this work shows metabolically labeled eASCs have the potential to be used for selective delivery of activity-inducing bioactive cues to enhance regenerative potential.

## 1. Introduction

Craniomaxillofacial (CMF) bone defects arise due to congenital abnormalities, oncologic resections, and high energy traumas. In 2023, around 470,000 people in US suffered from fractures of face bones or skull sustained from transport injuries or falls ^1^. These injuries heal poorly and are often large and irregularly shaped ^2^. The current standard of care relies on surgical reconstruction using autografts or allografts, but these methods face multiple challenges ^3–5^. Autograft approaches are limited by graft availability and can cause morbidity at the donor site, while allografts carry the risk of rejection and may perform poorly in high-risk settings (radiation, prior infection, age) ^5–7^. These limitations highlight the need for regenerative approaches that can restore bone while overcoming the constraints associated with traditional graft-based reconstruction. Regenerative biomaterials offer a promising strategy to address these challenges autografts and allografts ^4,8,9^. A wide range of synthetic and natural polymers as well as polymer-mineral composites have been described for broader bone tissue engineering applications. Our laboratory has recently described a mineralized collagen-glycosaminoglycan (MC) scaffold for craniofacial bone regeneration applications. We have optimized scaffold mechanics and mineral contents to accelerate osteogenic differentiation of bone marrow-derived mesenchymal stem cells (MSCs) or adipose-derived stem cells (ASCs) without the need for conventional osteogenic supplements (e.g., BMP-2) ^10–12^. We have also reported strategies to incorporate manuka honey or ascorbic acid to address microbial infection and boost cell activity, as well as incorporate 3D-printed polymeric mesh structures into the scaffold to aid conformal fitting and surgical handling ^8,13,14^. This acellular scaffold forms the basis of a translational work of bone regeneration.

Successful regeneration requires coordination of many processes such as both osteogenesis and angiogenesis at the implant-defect margin ^15,16^. While a range of biomaterials have been developed to support osteogenic activity of seeded or recruited osteoprogenitors such as mesenchymal or adipose derived stem cells, many fail to accelerate angiogenic infiltration ^16^. There is significant need to develop strategies to boost angiogenic-osteogenic coupling ^17^ without using supra-physiological doses of exogenous factors like BMP-2, BMP-7, and vascular endothelial growth factor (VEGF) ^3,18^. Osteoprogenitors such as MSCs and ASCs provide an avenue to address this challenge ^19^, They have the potential to act as endogenous producers of biomolecules, informed by the local biomaterial environment, to shape the trajectory of healing. Recently we showed modifications to MC scaffold structure and proteoglycan content can alter the secretome produced by resident MSCs or ASCs to increase regenerative potency ^20,21^. However, the complex nature of many craniofacial bone injuries, as well as the variability inherent in some complicated defects (e.g., post-radiation therapy; ageing; diabetes) ^17,22–25^ motivates the search for alternative strategies to boost angiogenic-osteogenic coupling. Our team has recently described a glycoengineering technique to achieve efficient and temporally controlled delivery of macromolecules *in vitro* and *in vivo*. Cells cultured in media containing tetraacetyl-*N*-azidoacetylmannosamine (Ac_4_ManAz) metabolize it to generate unique azide-functionalized glycoproteins and glycolipids on the cell membrane ^26–29^. These tags enable highly specific conjugation of macromolecules via efficient azide-dibenzocyclcooctyne (DBCO) click chemistry ^30–32^. Prior work showed conjugated macromolecules can be internalized to reach the cytosol and other organelles both in vitro and in vivo ^33–35^. Here, we will deploy this technology for the first time in a regenerative medicine application.

This work describes efforts to create engineered adipose-derived stem cells (eASCs) via metabolic glycan labeling followed by efforts to deliver VEGF to these eASCs within three-dimensional MC scaffolds. First, we report creation of engineered (Ac_4_ManAz-laden) ASCs and describe resultant azido tag density as a function of incubation time, sugar concentration, and as a function of cell proliferation. We then compare the regenerative activity of eASCs vs. non-engineered human ASCs (hASCs) within the MC scaffold using established metrics of cell activity (proliferation, metabolic activity) and gene expression. Finally, we report the degree to which eASCs can be actuated via the delivery of dibenzocyclooctyne (DBCO)-functionalized VEGF. We compare secretome (ELISA) and functional (Matrigel^TM^ vascular formation assay) metrics of actuated angiogenic potency for eASCs within MC scaffolds. We report a framework to tune the angiogenic potency of stem cells within the MC scaffold to accelerate efficient angiogenic and osteogenic activity necessary for regenerative healing.

## 2. Materials and Methods

This study first assessed the installation of Ac_4_ManAz metabolic glycan labels on human adipose stem cells. Engineered ASCs (eASCs) were then seeded onto mineralized collagen scaffolds to compare their regenerative potency to ASCs without labels. We subsequently compared the effect of selective delivery of supplemental VEGF functionalized with a DBCO tag on eASC angiogenic potency. ASC activity was assessed via gene expression profiles and VEGF secretion for up to 14 days, comparing results to the benchmark of human ASCs lacking metabolic tags (Figure 1).

**Fig. 1.**
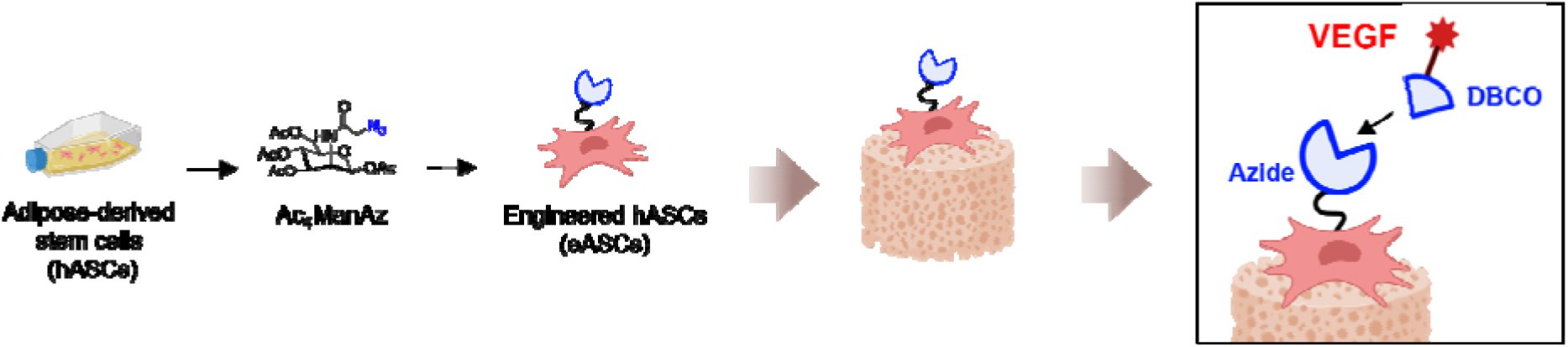
Schematic of experimental design to create engineered adipose stem cells (eASCs) via metabolic labeling, then assess regenerative potency and responsiveness to supplemental VEGF targeted to the installed azide moiety on eASCs.

### 2.1. Mineralized collagen-glycosaminoglycan scaffold fabrication

Mineralized collagen-glycosaminoglycan scaffolds were fabricated by homogenizing collagen type I from bovine (Collagen Matrix, Inc., Oakland, NJ, USA), chondroitin sulfate sodium salt (Spectrum Chemical MFG. Corp., Gardena, CA, USAs), calcium nitrate tetrahydrate (Sigma-Aldrich, St. Louis, MO, USA) and calcium hydroxide (Sigma-Aldrich) in 0.1456 M phosphoric acid (Sigma-Aldrich)/0.037 M calcium hydroxide buffer solution ^36–39^. The homogenized suspension was degassed and was pipetted into Aluminum 5” x 5” molds. The suspension was then a Genesis freeze dried (SP Scientific, Warminster, PA, USA) by decreasing the temperature from 20 °C to -10 °C at a constant rate of 1 °C/minute ^37^. The temperature was held at -10 °C for 2 hours to allow solidification, then sublimated at 0 °C and 0.2 Torr overnight ^37,39^. After lyophilization, fabricated collagen sheets were punched with a 6 mm biopsy punch (Acuderm Inc., Ft. Lauderdale, FL, USA) to form 6 mm diameter, 3 mm thick cylindrical specimens.

### 2.2. Scaffold sterilization, hydration, and crosslinking

Mineralized collagen scaffolds were sterilized for 12 hours with ethylene oxide using an AN74i Anprolene gas sterilizer (Andersen Sterilizers Inc., Haw River, NC, USA). Sterilized scaffolds were hydrated two days prior to cell seeding. Scaffolds were placed on the orbital shaker for moderate shaking throughout all incubation steps at room temperature. First, scaffolds were hydrated in ethanol for 90 minutes then washed with PBS for an hour. Scaffolds were crosslinked in EDC/NHS (EDC:NHS:COOH = 5:2:1) for 90 minutes followed by another PBS wash for an hour ^36,39^. Lastly, crosslinked scaffolds were placed in the cell culture medium (DMEM) for 48 hours at 37 °C prior to use, with media changed after 24 hours.

### 2.3. Synthesis of Ac_4_ManAz

₄ManAz was synthesized following a previously developed protocol . All chemical reagents for the synthesis were purchased from Sigma Aldrich unless otherwise noted. Mannosamine hydrochloride (1.0 equiv) was dissolved in anhydrous methanol and cooled to 0°C. Sodium methoxide in methanol (1.0 equiv) was then added, followed by chloroacetic anhydride (1.05 equiv) and triethylamine (1.0 equiv). The reaction mixture was stirred overnight at room temperature. After removing the solvent, the crude residue was redissolved in water, and sodium azide (4.0 equiv) was added. The mixture was heated at 60 °C overnight. The solvent was again removed, and the residue was dissolved in pyridine. Acetic anhydride and 4- dimethylaminopyridine were added, and the reaction was stirred for 24 h. The solvent evaporated, and the crude product was purified by silica gel column chromatography using ethyl acetate and hexane as eluents. 1H NMR of AAM (CDCl3, 500 MHz): δ (ppm) 6.66&6.60 (d, J= 9.0 Hz, 1H, C(O)NHCH), 6.04&6.04 (d, 1H, J=1.9 Hz, NHCHCHO), 5.32-5.35&5.04-5.07 (dd, J=10.2, 4.2 Hz, 1H, CH2CHCHCH), 5.22&5.16 (t, J=9.9 Hz, 1H, CH2CHCHCH), 4.60- 4.63&4.71-4.74 (m, 1H, NHCHCHO), 4.10-4.27 (m, 2H, CH2CHCHCH), 4.07 (m, 2H, C(O)CH2N3), 3.80-4.04 (m, 1H, CH2CHCHCH), 2.00-2.18 (s, 12H, CH3C(O)). 13C NMR (CDCl3, 500 MHz): δ (ppm) 170.7, 170.4, 170.3, 169.8, 168.6, 168.3, 167.5, 166.9, 91.5, 90.5, 73.6, 71.7, 70.5, 69.1, 65.3, 65.1, 62.0, 61.9, 52.8, 52.6, 49.9, 49.5, 21.1, 21.0, 21.0, 20.9, 20.9, 20.9, 20.8. ESI MS (m/z): calculated for C16H22N4O10Na [M+Na]+ 453.1234, found 453.1230.

### 2.4. Cell culture

#### Human adipose-derived stem cells

Human adipose-derived stem cells isolated from a 25-year-old Hispanic male with pre-existing conditions of hypertension, cirrhosis, and pancreatitis due to ethanol (hASCs; #205992; Essent Biologics, Centennial, CO, USA) were expanded from passage 4 to passage 5 in T175 flasks (ThermoFisher Scientific, Waltham, MA, USA) at 37 °C with 5% CO_2_ in RoosterBio expansion medium (RoosterBio, Frederick, MD, USA) for three days ^41^. After three days, media was changed to phenol-red free Dulbecco’s Modified Eagle Medium (DMEM) with low glucose and low glutamine (2nM) containing 10% fetal bovine serum (GeminiBio, West Sacramento, CA, USA), and 1% antibiotic-antimycotic (Gibco, Waltham, MA, USA) for two days.

#### Human umbilical vein endothelial cells

Human umbilical vein endothelial cells (HUVECs; Lonza, Walkersville, MD, USA) were expanded from passage 4 to passage 5 in T175 flasks at 37 °C with 5% CO_2_ in endothelial cell growth medium-2 (EGM-2, Lonza) for five days. Media was changed 24 hours after seeding and every two days from the first media change.

### 2.5. Scaffold seeding

hASCs or eASCs were seeded onto MC scaffolds at a density of 150,000 cells/scaffold via either static seeding or orbital seeding methods ^14,41–43^. For static seeding (for condition media exposure), 75,000 cells were pipetted onto one side of the scaffold in 24 µL of media for hASCs and 26 µL media for eASCs. Scaffolds were incubated for 30 minutes at 37 °C and 5% CO_2_ to allow initial attachment. Then, scaffolds were inverted and another 75,000 cells pipetted onto the other side of the scaffolds then incubated for an hour to allow further attachment. For orbital seeding (for functional and gene expression studies), scaffolds were placed in ultra-low attachment 6 well plates with hASCs (375,000 cells/mL). Scaffolds were incubated for 6 hours at 37 °C and 5% CO_2_ to allow for cell attachment. After seeding, cells were conventional cultured in low glucose DMEM, with media changed every three days.

### 2.6. DNA isolation and cell metabolic activity

DNA was extracted from cell-seeded scaffolds utilizing the DNeasy Blood & Tissue Kit (Qiagen, Hilden, Germany) using previously described methods ^44^. Briefly, scaffolds were quartered then incubated sequentially with kit buffers. The DNA concentration was measured by NanoDrop Lite Spectrophotometer (ThermoFisher Scientific) ^41,45^, with cell number per scaffold calculated using a simple linear regression model obtained from DNA extraction from known cell numbers at day 0.

Cell metabolic activity per scaffold was evaluated via alamarBlue viability assay (DAL1100, Invitrogen, Carlsbad, CA, USA) ^14,46^. Cell-seeded MC scaffolds were incubated in 10% alamarBlue assay solution in media for 90 minutes at 37 °C and 5% CO_2_ on the orbital shaker. Then, the solution was collected for fluorescence measurements with fluorescent spectrophotometer (Tecan Infinite F200 Pro, Männedorf, Switzerland). Fold change of metabolic activity was calculated from a linear calibration curve generated on day 0 from known cell concentrations. Metabolic activity was normalized to cell number at each timepoint.

### 2.7. RNA isolation and NanoString gene expression evaluation

RNA was extracted using TRIzol and RNA Clean & Concentrator Kits (Zymo Research, Irvine, CA, USA). Samples were obtained prior to scaffold seeding (day 0) as well as from cell-laden scaffolds at day 7 and 14 of culture ^47^. Scaffolds were minced with a sterile razor blade on days 7 or 14 vortexed and incubated in 1 mL of TriZol (Invitrogen) for 5 minutes ^41^. For day 0 control, cells were directly transferred into a centrifuge tube for TRIzol incubation. Then, 200 µL chloroform (Sigma-Aldrich) was added into the mixture for an additional 3 minutes ^41^, transferred to Phasemaker tubes (ThermoFisher Scientific) and centrifuged at 15,000 g for 15 minutes at 4 °C. Clear supernatant was transferred into spin columns for additional washing. Following manufacturer’s protocol, RNA was washed with ethanol, DNAse I, RNA prep buffer, RNA wash buffer and eluted in 15 µL DNase/RNase-free water. The RNA concentration and purity was measured with a NanoDrop Lite Spectrophotometer (ThermoFisher Scientific). Collected RNA was stored at -80 °C until analysis. Isolated RNA was analyzed via NanoString nCounter using a custom designed NanoString nCounter panel with 19 osteogenic genes, 13 immunomodulatory genes, 3 angiogenic genes and 3 housekeeping genes (Supplementary Table S1). nSolver Analysis software (NanoString Technologies Inc., Seattle, WA, USA) was used to calculate log2 fold change for gene expression normalized to both day 0 gene expression and to defined housekeeping gene (GAPDH, GUSB, OAZ1) ^20^. We also calculated log 2 fold change values normalized to hASC Day 0 gene expression using the same three housekeeping genes ^48^.

### 2.8. Quantification of calcium and phosphorous contents in mineralized collagen scaffolds

Mineralized collagen scaffolds were collected on Day 14 and analyzed by inductively coupled plasma mass spectrometry (ICP-MS) as previously described ^14,41^. Collected scaffolds were washed with PBS, fixed in 4% paraformaldehyde overnight at 4 °C and washed three times with PBS. Fixed scaffolds were then frozen at -80 °C overnight and lyophilized. Freeze-dried samples were digested in nitric acid and subsequently digested further in a CEM MARS6 microwave digestion system (CEM Microwave Technology Ltd., Matthews, NC, USA). The digested samples were diluted with deionized water and analyzed with an inductively coupled plasma-mass spectrometer (NexION 350D ICP-MS, PerkinElmer, Pittsfield, MA, USA).

### 2.9. Labeling of adipose-derived stem cells

To evaluate the role of Ac_4_ManAz dose on labeling efficacy, ASCs were maintained in culture media containing Ac_4_ManAz at concentrations ranging from 25 μM to 200 μM ^33^ for 48 hours. To evaluate the role of exposure time, ASCs were treated with Ac_4_ManAz for 24 hours, 48 hours or 72 hours at a defined Ac_4_ManAz concentration (50 μM). After labeling, cells were collected, washed with 2% FACS buffer solution (2% FBS solution in PBS), then incubated with Sulfo DBCO-Cy5 (233F0, 1:200; Lumiprobe, Hunt Valley, MD, USA) and Calcein AM (C1430, 1:5000; ThermoFisher Scientific) for 30 minutes. Cells were washed and analyzed via flow cytometry (Attune NxT; ThermoFisher Scientific) ^33,40^.

Long term retention of azido tags was then performed by culturing hASCs and eASCs (50 μM Ac_4_ManAz for 48 hours) in T75 flasks for 7 days. After 3 days and 7 days of culture, mean Cy5 fluorescence intensity (MFI) of ASCs was evaluated by flow cytometry (FACSymphony^TM^, BD Biosciences, San Jose, CA, USA) and FCS Express 6 (De Novo Software, Pasadena, CA, USA). Briefly, both groups were stained with Sulfo DBCO-Cy5 for 30 mins then LIVE/DEAD™ fixable violet dead cell stain (L34963, 1:1000, ThermoFisher Scientific) for 30 mins, with washes in 2% FACS buffer solution between each step. Cells were fixed by 4% paraformaldehyde for 15 minutes prior to analysis ^33,40^.

### 2.10. Staining of adipose-derived stem cells for confocal imaging

hASCs and eASCs were seeded on coverslips treated with poly-D-lysine (Gibco) and washed with 1% BSA. Cells were then incubated with Sulfo DBCO-Cy5 for 30 minutes. After incubation, cells were washed and fixed by incubating in 4% paraformaldehyde for 15 minutes. Cells were washed, then stained for Hoechst 33342 (H1399, 1.2:2000; Invitrogen) for 30 minutes. Cells were washed with 1% BSA, then imaged with a Zeiss LSM 900 Confocal Microscope (Zeiss, Oberkochen, Germany). Samples were imaged with 405 nm and 640 nm laser at 40x objective with 0.6% laser intensity and 700 V master gain. Images were post-processed with Fiji for colorization based on the assigned fluorescence channel with no additional adjustments ^49^.

### 2.11. DBCO functionalization of VEGF

Human VEGF-165 Recombinant Protein (100-20-50UG, ThermoFisher Scientific) was dissolved in 0.1 M sodium bicarbonate (NaHCO_₃_). Sulfo-DBCO-NHS (Vector Laboratories, Newark, CA, USA) was prepared at 100 mg/mL in DMSO. A 100-fold molar excess of Sulfo-DBCO-NHS was added to the VEGF solution, and the reaction was allowed to proceed for 1 hr at room temperature with gentle mixing. The reaction mixture was then transferred to a 3 kDa MWCO Amicon centrifugal filter (Sigma-Aldrich) and centrifuged to remove unreacted Sulfo-DBCO-NHS. The degree of labelling was quantified via UV-Vis spectroscopy by comparing the protein peak at 280 nm and the DBCO peak at 311nm (Supplementary Figure S5) ^40^. Following confirmation of successful conjugation, the DBCO-VEGF conjugate was resuspended in 0.1% BSA for storage at -80 °C until use ^40^.

### 2.12. Scaffold culture of human adipose-derived stem cells and engineered adipose-derived stem cells for media collection

hASCs or eASCs were seeded into mineralized collagen scaffolds at a density of 150,000/scaffolds using the described static seeding method. On day 1 after seeding, excess media was removed then cell seeded scaffolds were treated with a single dose of 0.1 μg/mL of VEGF or DBCO-VEGF for 30 minutes (effective dose chosen based on prior work ^50^) before conventional culture media was added again; and additional control group was hASC-seeded scaffolds that received no VEGF or DBCO-VEGF. Media was changed every three days (day 4 and day 7) until the experimental endpoint, with aliquots of conditioned media stored at -20 °C for later analysis.

### 2.13. ELISA of collected conditioned media

The concentration of VEGF in the conditioned media was quantified by a human VEGF DuoSet ELISA kit (DY293B, R&D Systems, Minneapolis, MN, USA) following manufacturer instructions, Day 4 and Day 7 samples were diluted 1:9 and 1:3, respectively, in reagent diluent (1% BSA, Sigma-Aldrich) prior to analysis. Briefly, plates were coated overnight with capture antibody and blocked for an hour. Conditioned media samples were added and incubated, followed by sequential incubation with detection antibody, streptavidin-horseradish peroxidase (Strep-HRP) and substrate solution, with washing performed between each step. 4 N of H_2_SO_4_ was added to stop the reaction and absorbance was measured by Tecan Infinite F200 pro. VEGF concentrations were calculated by fitting absorbance values to a standard curve and corrected with the initial dilution factors.

### 2.14. Matrigel tube formation assay

Cultured HUVECs were dispersed in 1:1 EGM-2 and conditioned (day 7) media collected from hASCs or eASCs exposed to either VEGF or DBCO-VEGF ^51^. HUVECs mixed with either basal hASCs media (DMEM) or EGM-2 and HUVECs mixed with conditioned media collected from hASCs were used as controls. 10,000 HUVECs in 150 µL of each media condition was seeded per well for Matrigel culture with 96-well Corning® Matrigel® Matrix well plate (356259, Corning, Corning, NY, USA) ^51,52^. HUVECs were incubated for 12 hours before brightfield imaging with LSM 900 (Zeiss). Analysis of the resultant networks was performed using the Angiogenesis Analyzer toolset for ImageJ (NIH, Bethesda, MD, USA) ^51,53^.

### 2.15. Statistical analysis

Data statistics for box plots are presented with interquartile range (25% - 75%) ± one standard deviation with labeled mean values. Graph plots are presented with mean ± 1 standard deviation from the mean. NanoString graph plots were made using ggplot package in RStudio 4.5.1, otherwise,plots were prepared in Microsoft Excel and OriginPro 2025. All statistical analysis was performed using RStudio 4.5.1. Grubbs test was used to screen for outliers. Then, the data set was tested for normality and variance with Shapiro-Wilk test and Levene’s test, respectively. The cutoff point for both tests was set to p < 0.05. Data with normal distribution and equal variance were tested by ANOVA and Turkey’s Honestly Significant Different (HSD) post-hoc test. Normal but not equal data was tested by Welch’s ANOVA and Games-Howell post-hoc test. Non-normal and equal data was tested by Kruskal-Wallis and Dunn’s test with Benjamini-Hochberg method. Non-normal and not equal data were tested by Welch’s Heteroscedastic F test with trimmed means and winsorized variance with Games-Howell post-hoc test.

## 3. Results

### 3.1. Human adipose stem cells can be metabolically labeled with azido groups

detected by conjugating with DBCO-Cy5 followed by flow cytometry analysis (Figure 2A). ASCs incubated in PBS (control) showed limited Cy5 signal, confirming limited non-specific binding of DBCO-Cy5 to unmodified ASCs. However, all tested concentrations of Ac_4_ManAz significantly increased the fraction of Cy5+ ASCs as well as the Cy5 mean fluorescence intensity (MFI) compared to control ASCs (p < 0.05). The percentage of Cy5^+^ cells rapidly approached 100%, with significant (p < 0.05) increases after exposure to 25 μM or 50 μM Ac_4_ManAz; (Figure 2B). Interestingly, the Cy5 MFI continued to increase significantly (p < 0.05) for each Ac_4_ManAz incubation density (Figure 2C). Tag density is directly proportional to Ac_4_ManAz concentration used during the incubation period.

**Fig. 2.**
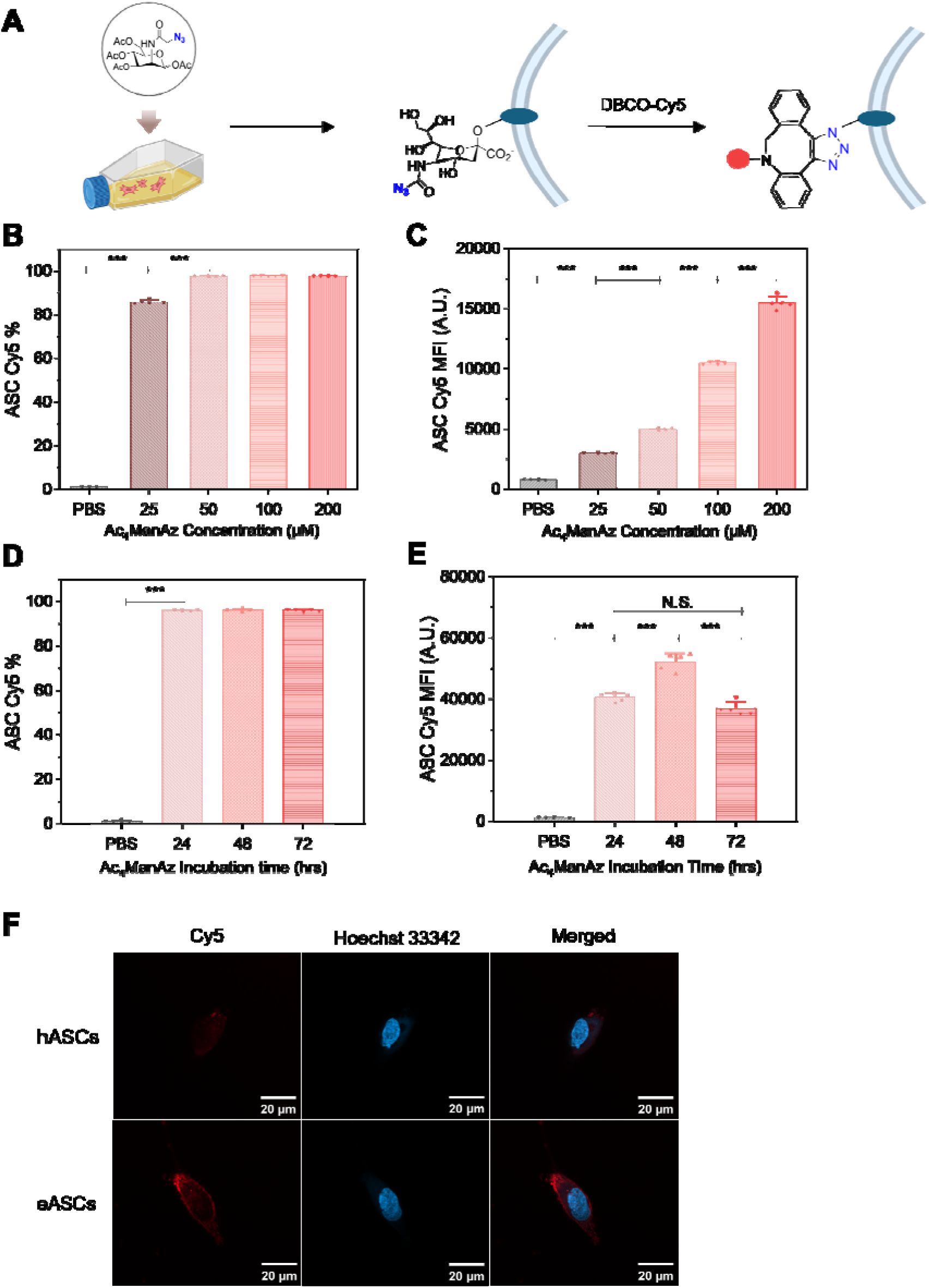
**A.** Schematic of metabolic glycan labeling of hASCs as well as analysis of label retention via DBCO-Cy5 probes. **B-C.** The percentage of Cy5^+^ eASCs and their Cy5 mean fluorescent intensity (MFI) increases with Ac4ManAz concentration (48 hour exposure; n=5). **D-E.** The percentage of Cy5^+^ eASCs and Cy5 MFI increases with Ac_4_ManAz incubation time (50 μM concentration; n=5). **F**. Control (hASC) and tagged (eASC) stained with sulfo-DBCO-Cy5 (red) and Hoechst 33342 (blue) after 48hour exposure to 50μM Ac_4_ManAz. N.S.: p > 0.05; *: p < 0.05; **: p < 0.01; ***: p < 0.001.

We subsequently defined the effect of Ac_4_ManAz incubation time using the Ac_4_ManAz concentration (50 μM) found to efficiently label ASCs with excellent cell viability (Figure 2D-E). We observed efficient Ac_4_ManAz labeling, with significant (p < 0.05) increase in the fraction of Cy5^+^ ASCs after 24 hours of Ac_4_ManAz exposure, with near 100% labeling, and no significant change (increase or decrease) in labeling for increasing Ac_4_ManAz exposure time (48, 72 hours; Figure 2D). While the fraction of labeled ASCs remains unchanged after 24 hours exposure, the Cy5 MFI showed further significant increase up to 48 hours of exposure, suggesting increasing numbers of azido tags per cell with increased incubation time. hASCs incubated for 72 hours in Ac_4_ManAz showed a decrease in Cy5 MFI compared to cells incubated for 48 hours (p < 0.05; Figure 2E), which can be attributed to the dilution effect over cell divisions. To visualize azido labeling, hASCs (control: PBS incubation) and eASCs (50 μM of Ac_4_ManAz for 48 hours) were subsequently stained with Sulfo-DBCO-Cy5 (to label the azido tag) and Hoechst 33342 (to label all nuclei) on Day 3 (72 hours after washing out Ac_4_ManAz) then imaged with confocal microscopy. eASCs showed stronger Cy5 fluorescence localized on the cell membrane compared to hASCs (Figure 2F).

### 3.2 eASCs retain Ac_4_ManAz tags through 7 days of in vitro culture

After establishing a framework to install azido tags on hASCs to create a population of eASCs, we subsequently evaluated azido tag retention (Figure 3). eASCs were generated by exposing hASCs to 50 μM of Ac_4_ManAz for 48 hours; hASCs and eASCs were then separately cultured for 7 days in two-dimensional culture. Populations of hASCs and eASCs both expanded over 7 days in culture. By day 7 the number of (non-modified) hASCs was significantly (p < 0.05) greater than eASCs (Figure 3A). The vast majority of eASCs continued to contain azido tags (Cy5-DBCO label) through 3 days in culture (Fig. 3B). While after 7 days in culture the fraction of eASCs retaining azido tags was reduced, expected given multiple cell doublings that occurred in culture, eASCs still exhibited significantly (p > 0.05) greater azido tags than non-modified hASCs (more than 25% still containing azido tags). Similar trends were observed for the Cy5 MFI for eASCs after 3 and 7 days, with significantly (p < 0.05) greater tags on eASCs than hASCs (Figure 3C).

**Fig. 3.**
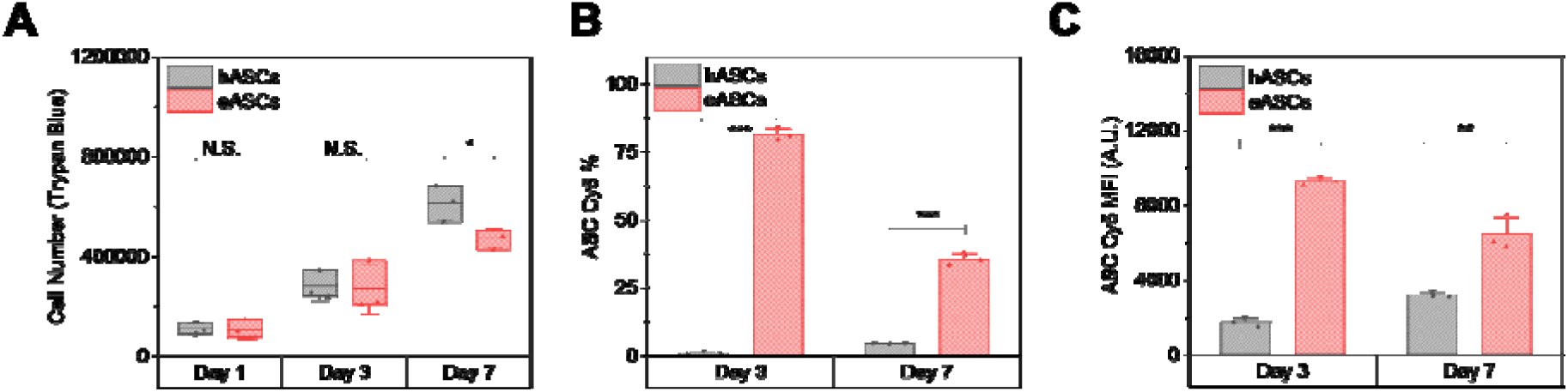
hASCs and eASCs each cultured in T75 flasks for 7 days. **A**. Number of eASCs and hASCs with time (Trypan Blue; n=3). **B-C**. The percentage of Cy5^+^ eASCs and Cy5 MFI after in vitro culture (n=3). N.S.: p > 0.05; *: p < 0.05; **: p < 0.01; ***: p < 0.001.

### 3.3 Azido-labeled eASCs can proliferate and are metabolically active within three-dimensional mineralized collagen scaffolds

We then examined if eASCs display distinct expansion and activity metrics compared to non- modified hASCs when cultured within 3D mineralized collagen scaffolds. eASCs were generated by exposing hASCs to 50 μM Ac_4_ManAz for 48 hours prior to scaffold seeding.

hASCs or eASCs were seeded onto mineralized collagen scaffolds and cultured for 14 days Both hASCs and eASCs displayed identical expansion metrics, with ∼40% of seeded cells adhering 1 day after seeding then cell proliferation being observed after 7 days in culture (p > 0.05; Figure 4A). Both hASCs and eASCs also showed increased metabolic activity with culture time. Notably, eASCs containing azido tags were significantly (p < 0.05) more metabolically active than non-modified hASCs for all time points (Figure 4B).

**Fig. 4.**
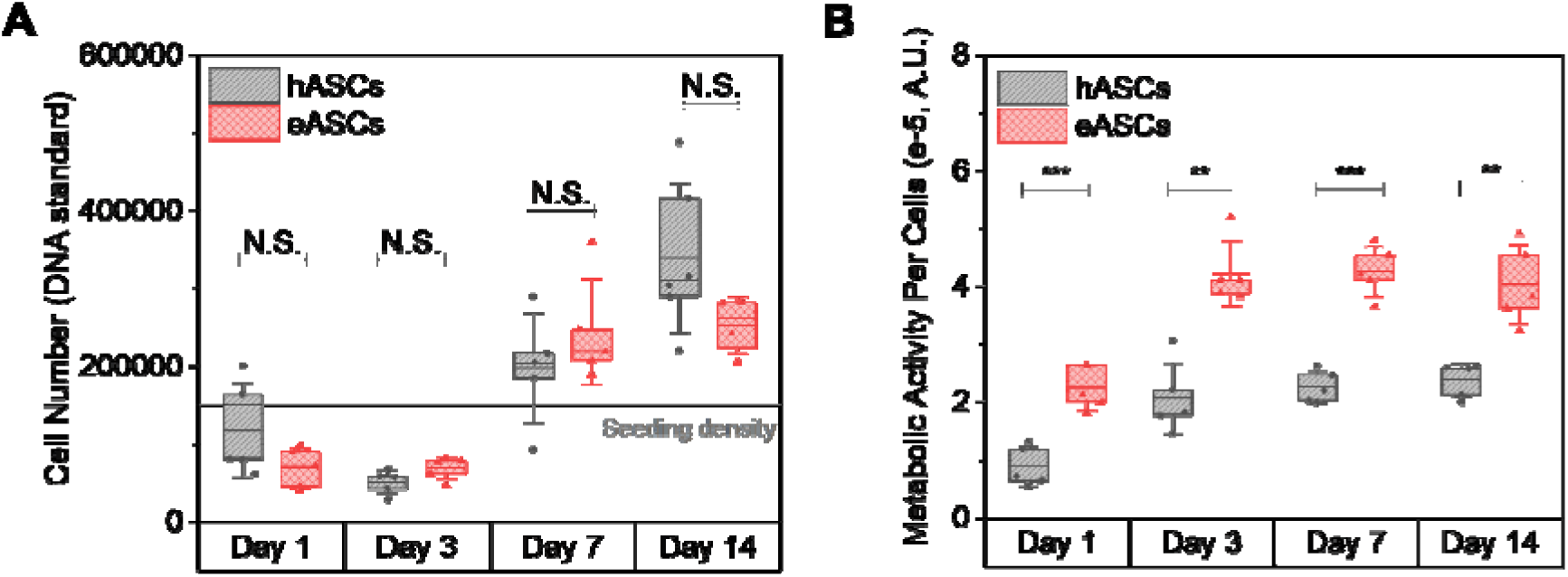
hASCs and eASCs viability across timepoints. **A.** Quantifying numbers of hASCs and eASCs cultured in mineralized collagen scaffolds for 14 days. **B.** Metabolic activity per cells for eASCs and hASCs in mineralized collagen scaffolds as a function of culture time. N.S.: p > 0.05; *: p < 0.05; **: p < 0.01; ***: p < 0.001.

### 3.4 Azido-tagging does not reduce osteogenic, angiogenic, or immunomodulatory potency in mineralized collagen scaffolds

We then examined if the addition of azido tags influenced the regenerative potency of ASCs within the mineralized scaffolds, using analysis of a library of osteogenic, angiogenic, and immunomodulatory genes previously developed to benchmark mineralized collagen scaffold potency using (non-modified) hASCs. Gene expression levels were measured using NanoString and reported as log 2 fold change normalized to hASCs respective Day 0 gene expression level (Figure 5, Supplementary Figure S2). Broadly, eASCs displayed remarkably similar angiogenic, osteogenic, and immunomodulatory gene expression levels and trajectories when compared to hASCs. There were some notable changes, with increased expression of osteogenic genes OPN and TNFSF11 and reduced levels of ALPL, RUNX2, and WNT16 in eASCs at Day 0 (p <0.05, respectively; Figure 5). We also observed significantly increased levels of the osteogenic gene TNFRSF11B in eASCs (p < 0.05, Figure S2) at days 7 and 14 of culture. We also observed some shifts in immunomodulatory genes, with IL8 and TSG6 expressed higher (p < 0.05), and CCL2 expressed significantly lower (p < 0.05) by eASCs, but only at day 0. We did not observe any difference in angiogenic gene expression.

**Fig. 5.**
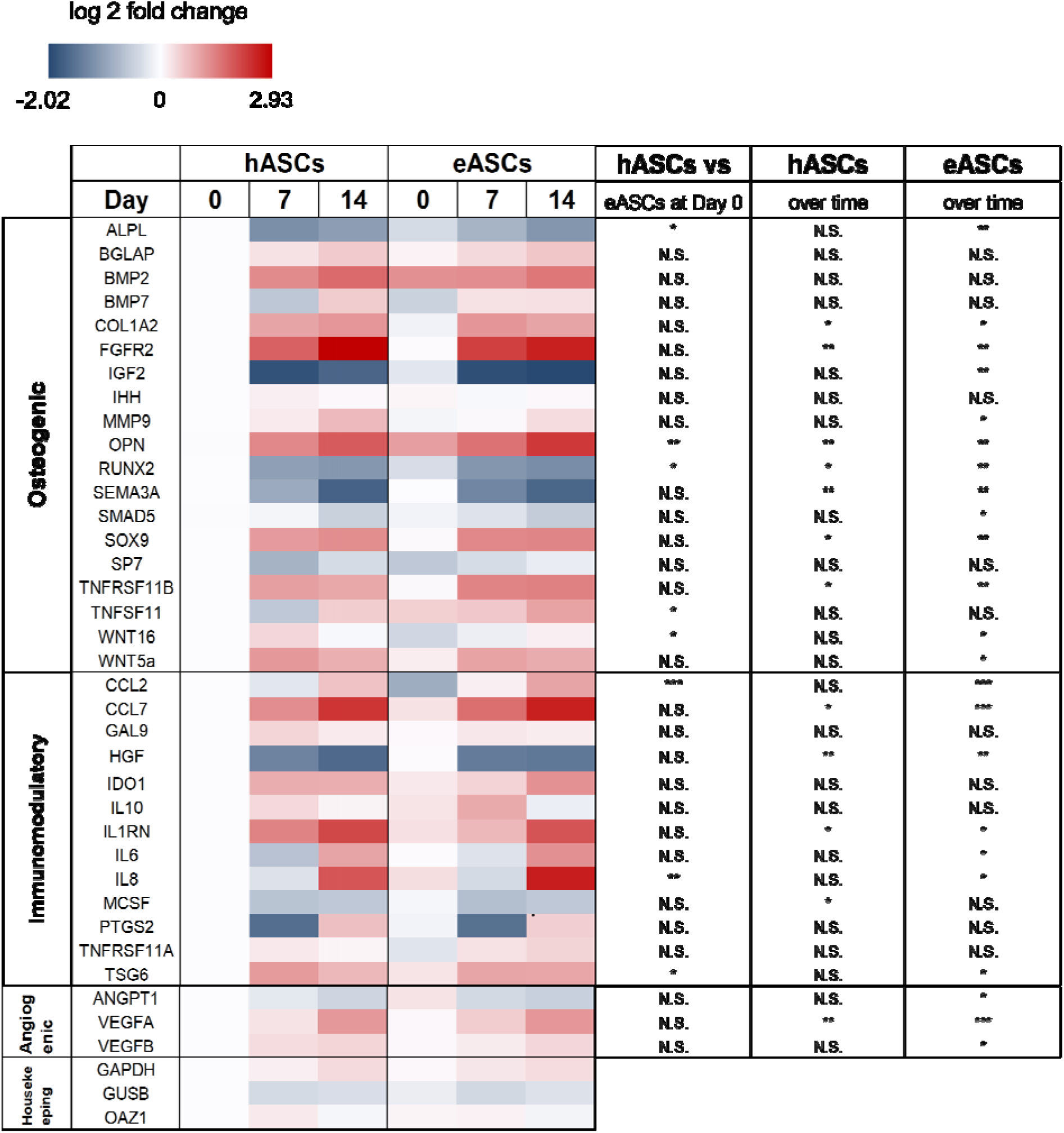
Gene expression levels for hASCs and eASCs cultured for up to 14 days in mineralized collagen scaffolds measured via a custom NanoString panel. Expression levels are reported as log 2 fold change normalized to initial expression by hASCs at day 0 prior to seeding and to established housekeeping genes (GAPDH, GUSB, OAZ1). Statistical comparisons were made for: initial (day 0) differences in hASC vs. eASC gene expression; significant changes in eASC or hASC expression trajectories for each gene with time. N.S.: p > 0.05; *: p < 0.05; **: p < 0.01; ***: p < 0.001.

We then examined trends of gene expression changes in hASCs and eASCs over the culture period via a linear trend test (Figure 5). We observed consistent patterns of upregulation of COL1A2, FGFR2, OPN, SOX9, and TNFSF11 and downregulation of RUNX2 and SEMA3A with time (day 0 through 14) for both hASCs and eASCs (P < 0.05). We also observed upregulation of MMP9, WNT16, and WNT5a and downregulation of ALPL, IGF2, and SMAD5 with time in eASCs only (p < 0.05). Examining immunomodulatory genes, we observed consistent upregulation of CCL7 and IL1RN and downregulation of HGF for both hASCs and eASCs in mineralized collagen scaffolds (p < 0.05). We observed reduced expression of MCSF with time in only hASCs, and upregulation of CCL2, IL6, IL8, and TSG6 in only eASCs (p < 0.05). And considering angiogenic genes, we observe upregulation of VEGFA for both hASCs and eASCs, while upregulation of VEGFB and downregulation of ANGPT1 was more pronounced in only eASCs.

### 3.5 DBCO-VEGF can be internalized by eASCs

We evaluated the functional consequence of the addition of DBCO-VEGF, which can conjugate to azido-tagged cells, into hASC and eASC seeded mineralized collagen scaffolds. Conditioned media was collected for 7 days from hASC and eASC seeded mineralized collagen scaffolds. On Day 1, 0.1 μg/mL of VEGF (control) or 0.1 μg/mL of DBCO-VEGF (clickable to azido group) was administered onto hASC or eASC seeded mineralized collagen scaffolds (Figure 6A). Cumulative VEGF produced over 7 days was quantified via ELISA (Figure 6B, Supplementary Figure S5). No significant difference in VEGF in the conditioned media was observed for hASCs or eASCs at Day 4. However, conditioned media collected from eASCs exposed to DBCO- VEGF (eDV) contained significantly greater amounts of VEGF at Day 7 compared to hASCs exposed to DBCO-VEGF (hDV) (p< 0.05). No significant increase was observed for hASCs (hDV group) between Day 4 and 7 for the DBCO-VEGF treated group (p > 0.05). We also did not observe any difference in cumulative VEGF for hASCs or eASCs treated with conventional (not Ac_4_ManAz targeted DBCO-VEGF) VEGF (hV and eV groups; p > 0. 05). Lastly, we examined if the levels of VEGF produced by ASCs were sufficient to induce functional changes in HUVECs via a Matrigel tube formation assay. Briefly, HUVECs were cultured for 12 hours in conditioned media collected from each ASC group after 7 days (hDV, eDV, hV, eV). However, we did not observe any differences in HUVEC network formation among the groups (Figure 6C, Supplementary Figure S6).

**Fig. 6.**
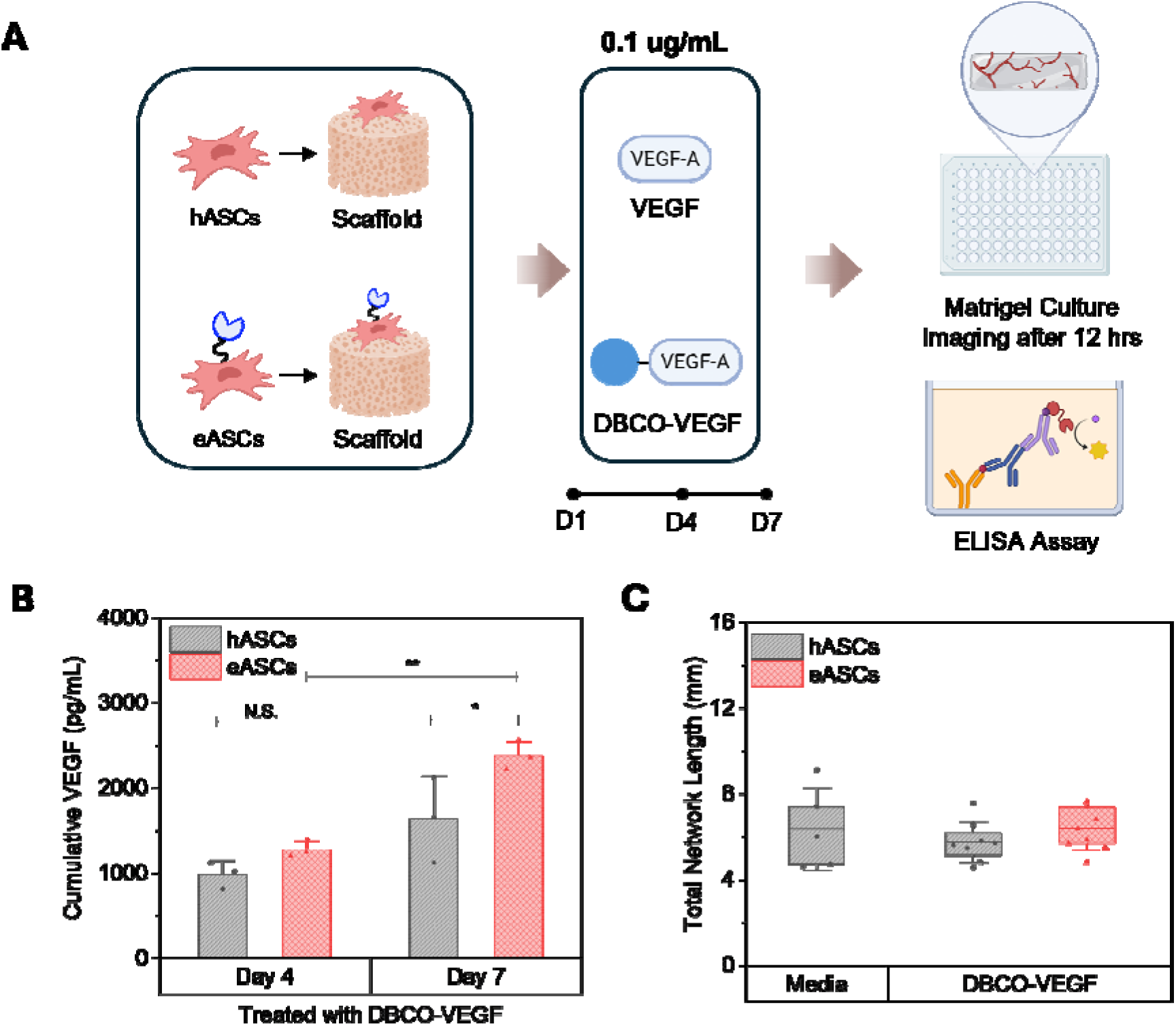
Tagging actuation with DBCO-VEGF. **A.** Schematic of the analysis of eASC vs. hASC response to DBCO-tagged VEGF. **B.** Cumulative VEGF secreted into culture media by eASCs vs. hASCs treated with DBCO-VEGF and cultured in mineralized collagen scaffolds, measured via ELISA. **C.** Total length (mm) of HUVEC networks formed in Matrigel culture in response to base (Media) or conditioned media generated by DBCO-VEGF stimulated hASCs or eASCs in mineralized collagen scaffolds. N.S.: p > 0.05; *: p < 0.05; **: p < 0.01.

## 4. Discussion

Osseous defects of the skull occur secondary to trauma, congenital abnormalities, or after resection to treat stroke, cerebral aneurysms, or cancer ^54–59^. A primary focus for regenerative strategies to promote craniofacial reconstruction has been to accelerate osteogenic differentiation of osteoprogenitors such as mesenchymal stem cells or adipose stem cells. This has included the delivery of osteogenic factors such as bone morphogenetic protein (BMP-2) ^60^; however, the typical need for supraphysiological doses have led to significant complications such as ectopic bone formation, resorption of adjacent bone, and maxillary growth disorders ^61–65^. More recently, efforts have turned to the design of biomaterials that provide structural, compositional, and mechanical signals that accelerate osteogenic differentiation in the absence of exogenous cytokines. Via a series of mechanistic investigations, we identified a mineralized collagen (MC) scaffold that endogenously activates BMP receptor (BMPR) signaling and promotes expansion and osteogenic differentiation of MSCs and ASCs without exogenous BMP ^44,63,66–68^. This scaffold now provides the basis for the seeding or recruitment of osteoprogenitor cells within the wound site as well as design features which accelerate osteogenic activity. However, impaired angiogenesis at the defect-implant margin is a predisposing factor for failed healing ^17,69–72^. And further, the degree of angiogenic response can be significantly influenced by the scale of defect as well as a wide range of patient-specific factors including age and underlying disease.

Hence the objective of this project was to advance strategies that accelerate angiogenic-osteogenic coupling within the scaffold using hASCs augmented with metabolic glycoengineering motifs to exogenously direct actuation of their angiogenic phenotype *in situ*. We described strategies to create engineered ACSs (eASCs) that can be seeded into a porous mineralized collagen biomaterial. While metabolic glycoengineering have been used to install label on a range of early progenitor (e.g., induced pluripotent stem cells, embryonic stem cells) and immune cells ^26,33,34,40,73^, their utility in a regenerative setting using adipose stem cells have not been previously shown. We report titratable control over the installation of azido groups onto hASCs as a function of Ac_4_ManAz concentration and exposure time. We identify an optimal dosing regimen (50 μM incubation for 48 hours) and confirm azido-tagged eASCs are proliferative, metabolically active, and with high azido tag retention for at least 3 days tag, while a significant minority retain tags for at least 7 days in two-dimensional culture. As a result, we identify a therapeutic window (< 7 days) for the delivery of DBCO-macromolecules to eASCs within the scaffold. This window is largely consistent with prior work using azido-tagged immune cells ^33,73^, suggesting a broader use for selectively tagged cells for regenerative medicine applications. This 7-day window is also highly relevant for our specific application to boost implant angiogenic potency given that the first two weeks after implant are critical for vascular healing, angiogenic-osteogenic coupling, and eventual regeneration ^74–82^. One area for future exploration based on this project is an improved understanding of tag retention within fully three-dimensional biomaterials. A primary route for the loss of membrane tags is cell division, where installed tags are split between two daughter cells. While we show eASCs retain tags for up to 7 days in two-dimensional culture, ASC proliferation is often slower in fully three-dimensional tissue engineering biomaterials. As a result, eASCs may retain metabolic tags far longer in fully three-dimensional biomaterials. However, methods to recover ASCs from 3D scaffolds (e.g. enzymatic degradation) may also affect tag retention, making such studies significantly more difficult.

This project subsequently validated that the installation of azido tags did not significantly alter the baseline regenerative potency of eASCs via analysis of a panel of osteogenic, immunomodulatory, and angiogenic genes for hASCs and eASCs seeded in mineralized collagen scaffolds for up to 14 days of culture. This panel of genes was previously developed during the optimization of the mineralized collagen scaffold ^44,83^, with findings here also largely consistent with prior work developed metabolically tagged immune cells ^35,73^. And lastly, we showed delivery of the angiogenic factor VEGF in a DBCO-tagged format selectively increased VEGF secretion of eASCs (vs. hASCs) during the identified 7-day window. Notably, this effect was pronounced in eASCs only for DBCO-tagged VEGF, and not for conventional soluble VEGF that worked equivalently on eASCs and hASCs. While all experiments here were in vitro, prior work has also demonstrated the increased accumulation of DBCO-tagged payloads in metabolically labeled cell in vivo ^40^, suggesting future efforts to implant then actuate eASC- seeded scaffolds in vivo based on the kinetics of angiogenic healing. This approach also suggests future efforts to consider the use of a small fraction of eASCs within a large non-modified ASC population to shape healing kinetics, as well as the use of multiple orthogonal metabolic tags to selectively activate multiple regenerative trajectories independently.

## 5. Conclusions

Craniofacial bone regeneration is a complex process involving the coordinated action of progenitor cells to promote osteogenic and angiogenic trajectories. While the field of bone tissue engineering has largely concentrated on biomaterial designs to induce progenitor differentiation towards osteogenic phenotypes, impaired angiogenic response can significantly reduce the quality of healing. As a result, there is an opportunity to consider strategies that orchestrate angiogenic-osteogenic coupling. Using a mineralized collagen scaffold recently identified to promote osteogenic activity of ASCs in the absence of exogenous osteogenic supplements, here we report the use of metabolic glycoengineering methods to install azido tags into ASCs to create an engineered ASC (eASC) population. We confirm processing conditions to enable efficient azido tag installation as well as retention for up to a week in vitro. Installation of azido tags does not alter eASC osteogenic, angiogenic, and immunomodulatory phenotypes within the mineralized collagen scaffold. However, using DBCO-azide click chemistry, we show exposure to DBCO-tagged VEGF increased the angiogenic activity of eASCs versus non-modified ASCs. This study suggests the potential for employing populations of eASCs within mineralized collagen scaffolds to shape the angiogenic phenotype via orthogonal, exogenous signals to enhance regenerative potency, establishing the basis of coordinated use of engineered biomaterials and synthetically engineered stem cells to improve regenerative outcomes.

## Acknowledgements

The authors would like to acknowledge the following institutes for access to their facilities and services: the Carl R. Woese Institute for Genomic Biology, the Cytometry and Microscopy to Omics Facility, and the School of Chemical Sciences Microanalysis Laboratory, located at the University of Illinois. This manuscript is the result of funding of the National Institute of Dental and Craniofacial Research of the National Institutes of Health under Award Number R01 DE030491 (BACH). It is subject to the NIH Public Access Policy. Through acceptance of this federal funding, NIH has been given a right to make this manuscript publicly available in PubMed Central upon the Official Date of Publication, as defined by NIH. Imaging and transcriptomic analyses were performed using the Tumor Engineering and Phenotyping, the Carver Biotechnology Center, and the Multimodal Biomedical Imaging Shared Resources within the Cancer Center at Illinois (CCIL), which is partially supported by the NCI Cancer Center Support Grant (CCSG) award number P30CA275774. The content is solely the responsibility of the authors and does not necessarily represent the official views of the National Institutes of Health. Additional support was provided by the Carl R. Woese Institute for Genomic Biology, the Dept of Chemical and Biomolecular Engineering, and the Dept. of Materials Science and Engineering at the University of Illinois at Urbana-Champaign.

## Contributions (CRediT: Contributor Roles Taxonomy ^84,85^

**Jaejung Kim:** Conceptualization, Data curation, Formal Analysis, Visualization, Investigation, Methodology, Writing – original draft, Writing – review & editing. **Dhyanesh Baskaran:** Conceptualization, Investigation, Data curation, Methodology, Formal Analysis, Writing – review & editing. **Alison Nunes:** Investigation. **Génesis Ríos Adorno:** Investigation. **Hua Wang:** Supervision, Conceptualization, Resources, Funding acquisition, Methodology, Writing – review & editing. **Brendan A.C. Harley:** Conceptualization, Resources, Project administration, Funding acquisition, Supervision, Writing – review & editing.

## Data Availability

The data that support the findings of this study are available on request from the corresponding author.

