## Supplemental Information for "Metabolic tagging of adipose-derived stem cells for targeted modulation of regenerative potency within mineralized collagen scaffolds"

600 S. Mathews Ave.

Urbana, IL 61801

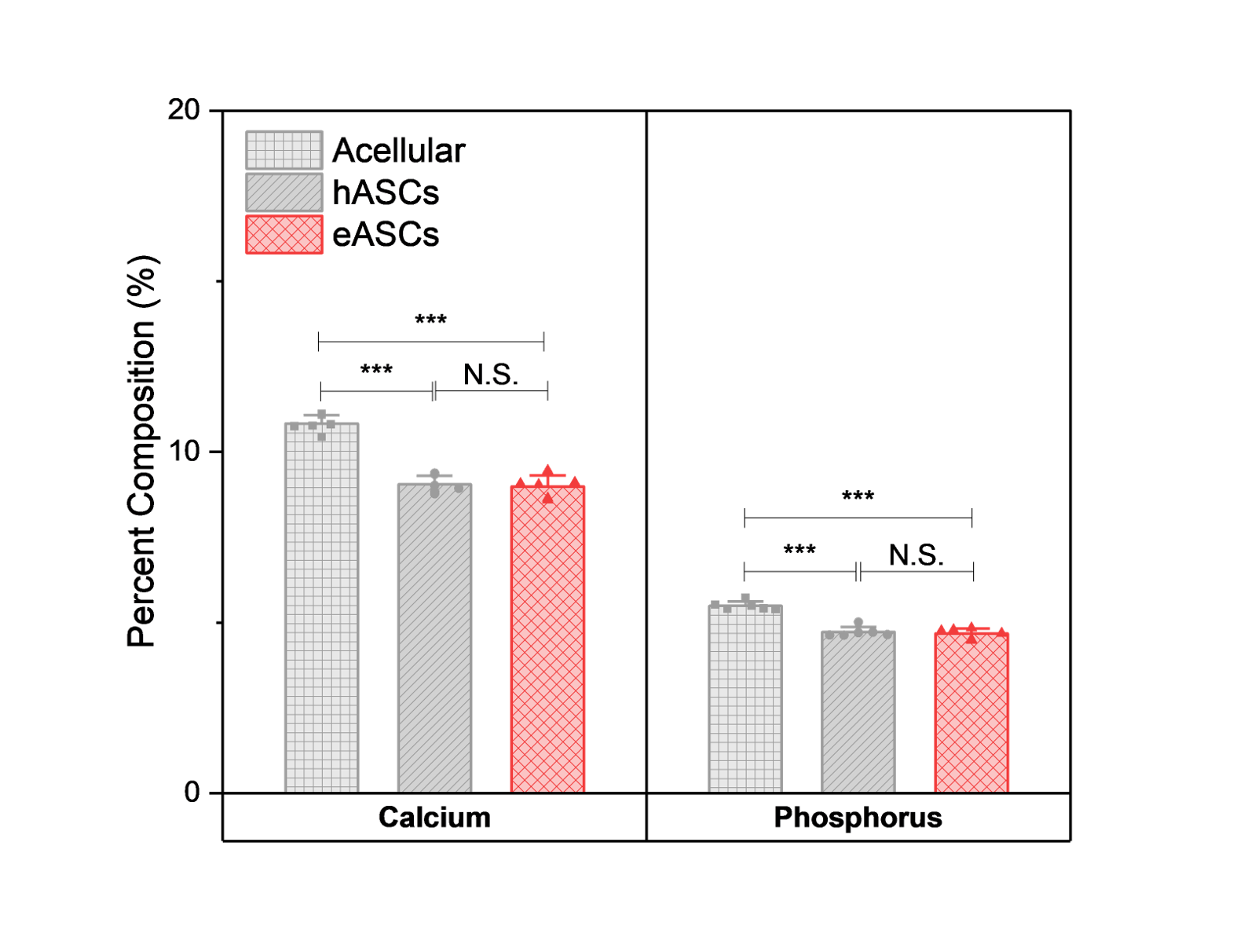

Figure S1. hASC/eASC scaffold mineral remodeling. hASCs and eASCs were cultured in mineralized collagen scaffolds for 14 days. After culture, scaffolds were fixed and overall mineral composition was evaluated by Inductively coupled plasma mass spectrometry (ICP-MS). N.S.: p > 0.05; *: p < 0.05; **: p < 0.01; ***: p < 0.001.

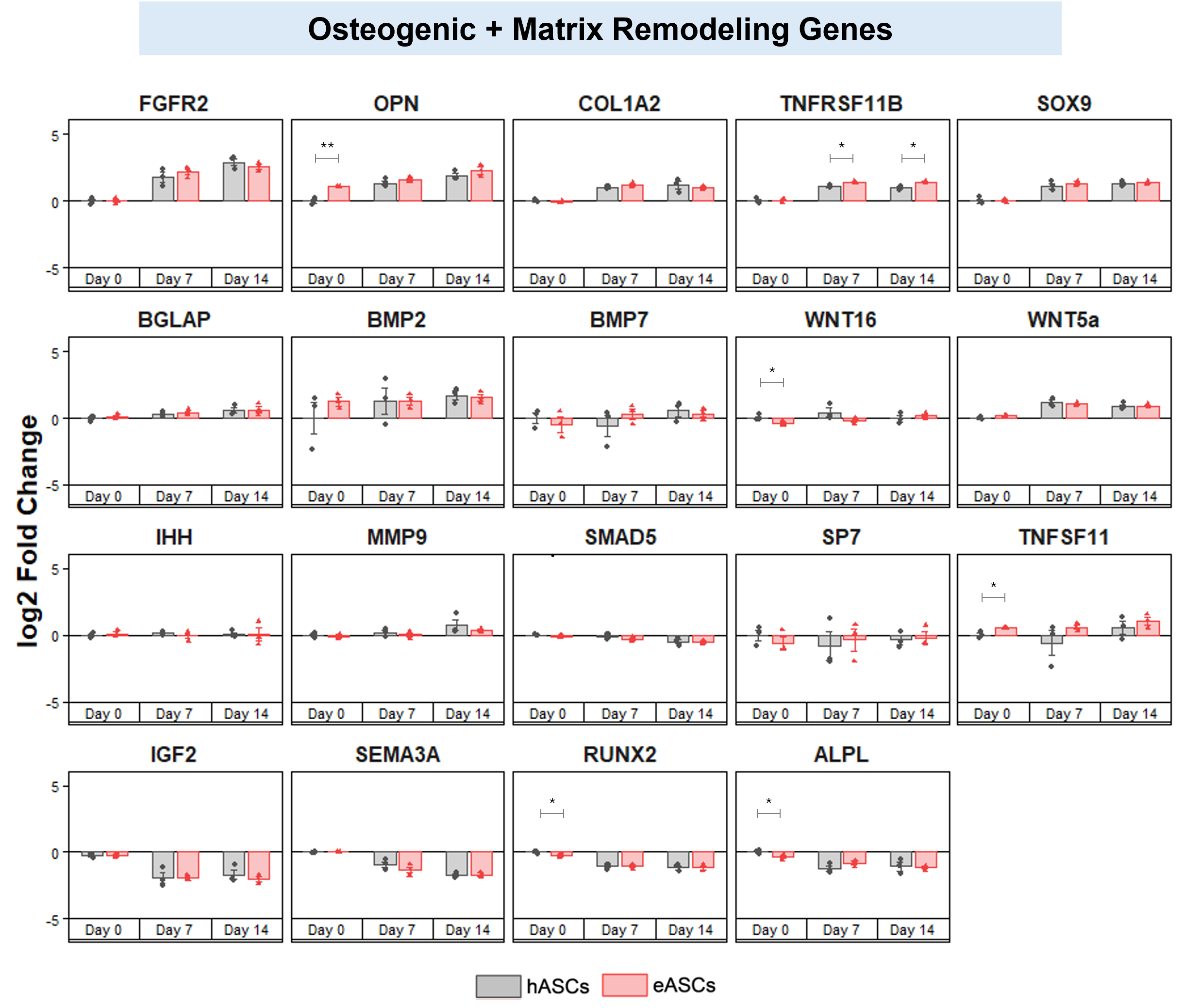

Figure S2. Osteogenic trajectories. Fold change of a panel of osteogenic genes for hASCs and eASCs cultured for up to 14 days in mineralized collagen scaffolds via a custom NanoString panel. N.S.: p > 0.05; *: p < 0.05; **: p < 0.01; ***: p < 0.001.

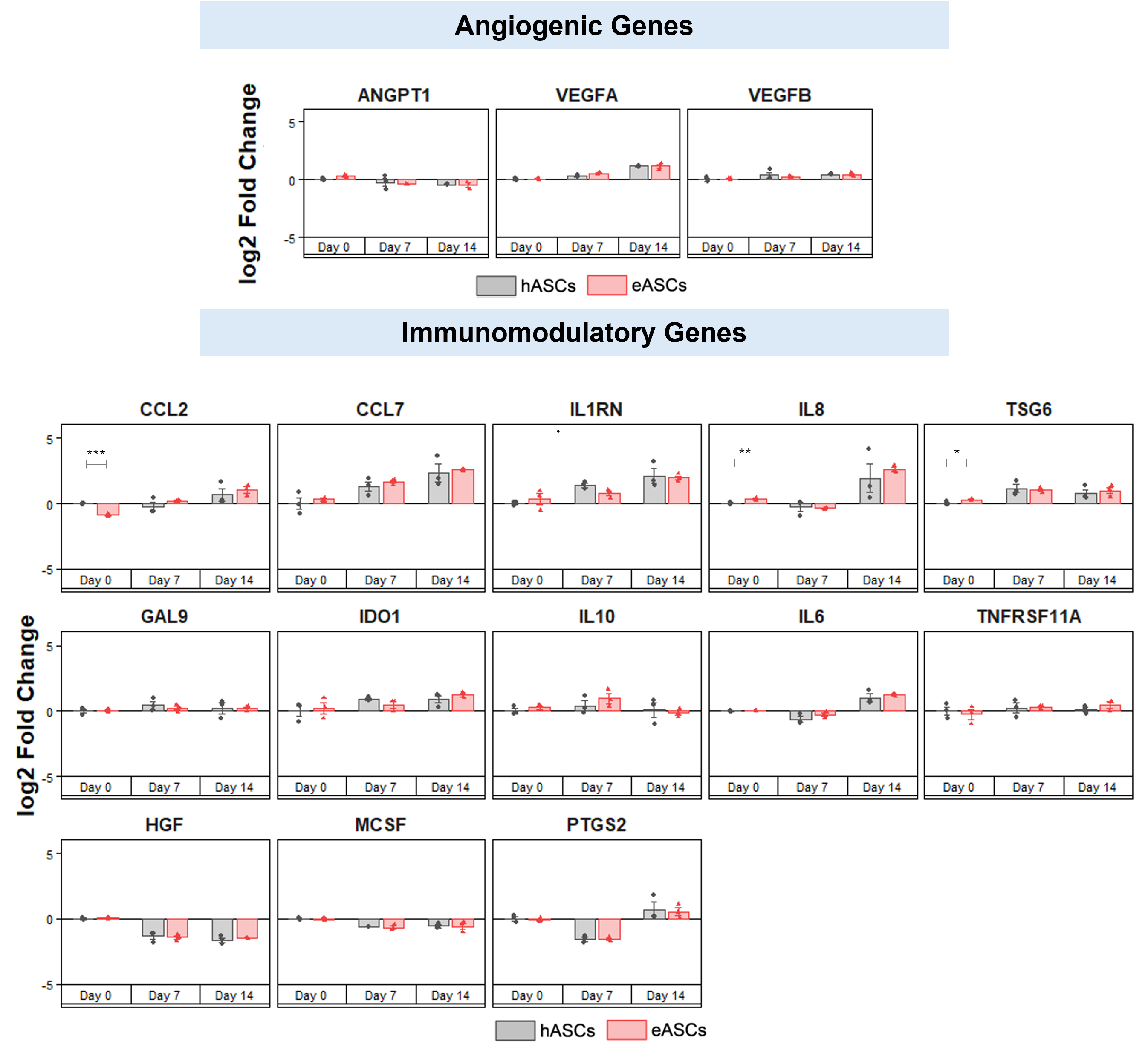

Figure S3. Angiogenic and immunomodulatory trajectories. Fold change of a panel of angiogenic and immunomodulatory genes for hASCs and eASCs cultured for up to 14 days in mineralized collagen scaffolds via a custom NanoString panel. N.S.: p > 0.05; *: p < 0.05; **: p < 0.01; ***: p < 0.001.

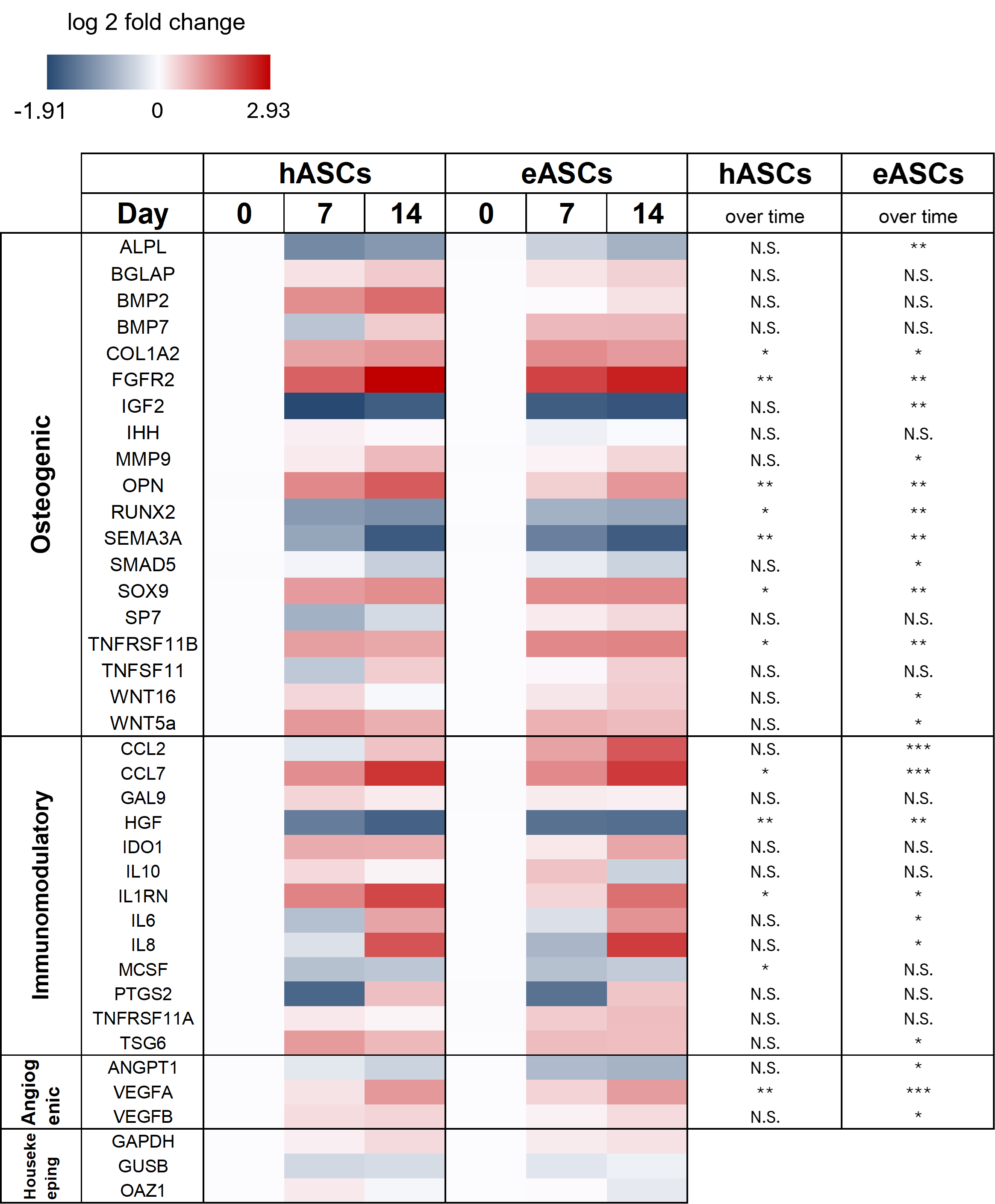

Figure S4. Heat map representing relative phenotype gene expression trajectories. Log 2 fold change of hASC and eASC gene expression over 14 days in mineralized collagen scaffolds measured via a custom NanoString panel. Gene expression levels are log 2 fold change normalized to each group’s respective Day 0 expression level and to housekeeping genes (GAPDH, GUSB, OAZ1). Red indicates upregulation and navy indicates downregulation. The significance of overall up or downregulation of each gene with time over the culture period was also evaluated. N.S.: p > 0.05; *: p < 0.05; **: p < 0.01; ***: p < 0.001.

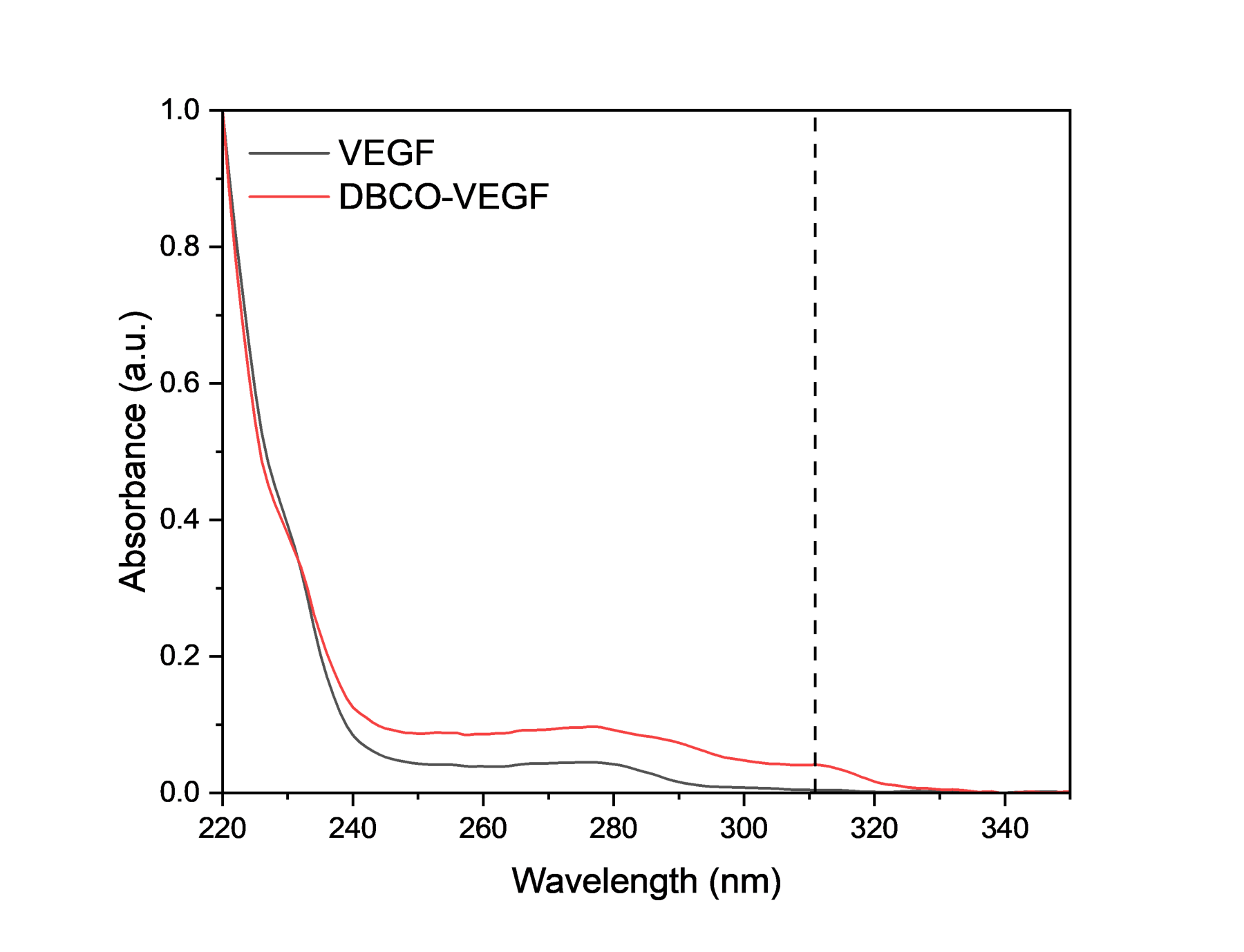

Figure S5. VEGF-165 was functionalized with DBCO group. The peak at 311 nm indicates successful functionalization of VEGF-165 with DBCO.

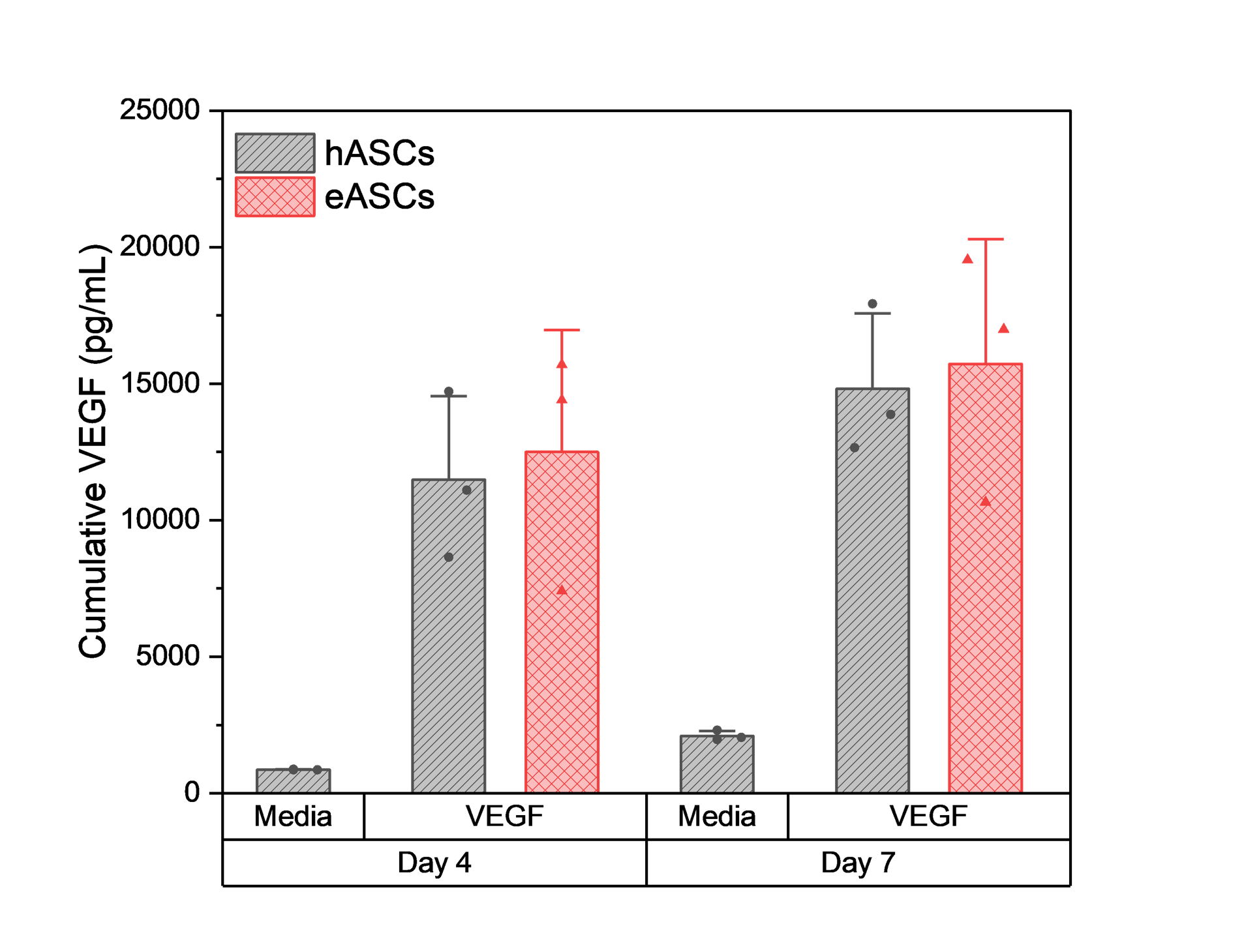

Figure S6. hASCs or eASCs were cultured in mineralized collagen scaffolds with basal media (DMEM) or human VEGF-165 for 7 days. Cumulative VEGF was measured via ELISA. N.S.: p > 0.05; *: p < 0.05; **: p < 0.01; ***: p < 0.001.

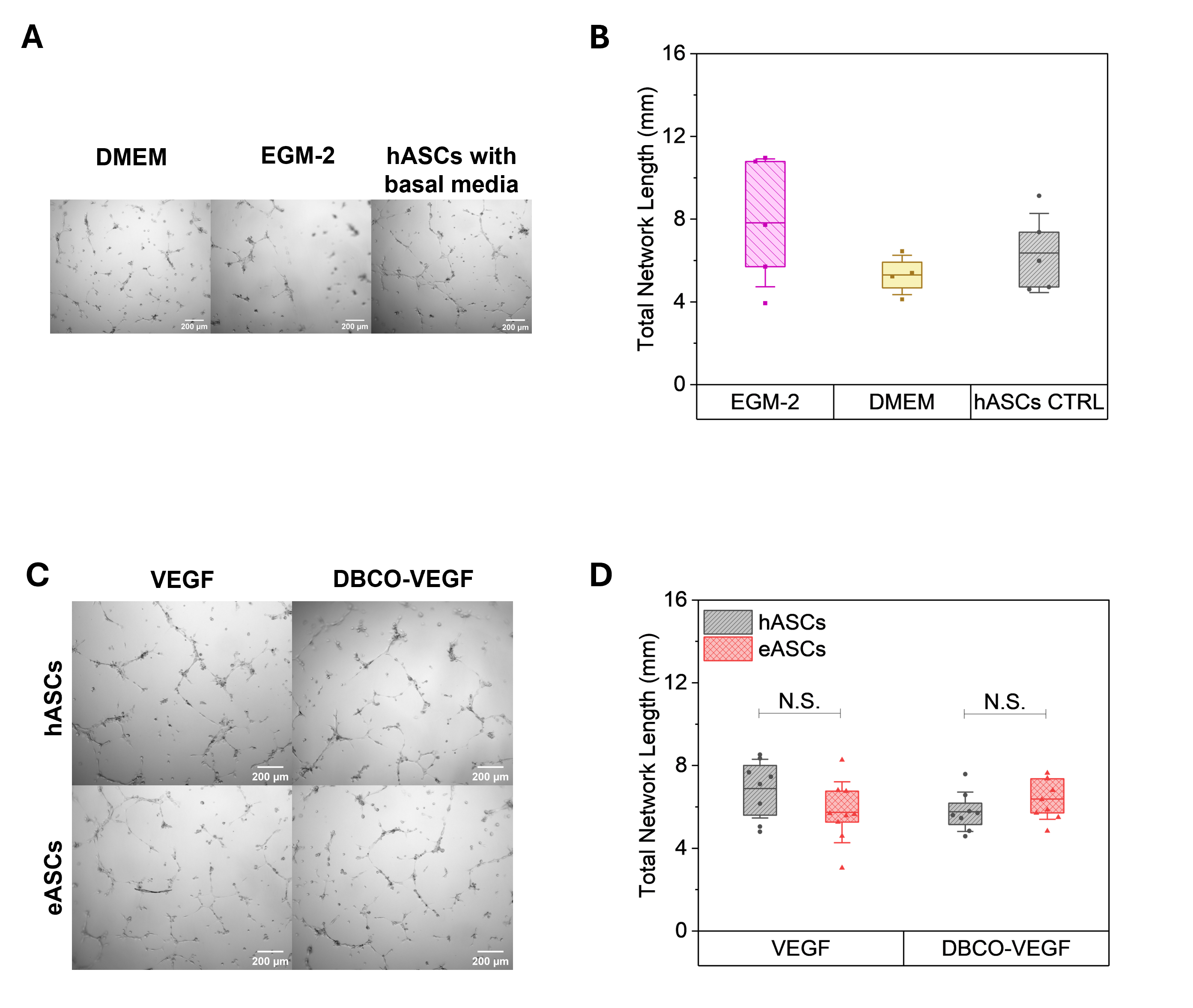

Figure S7. 12 hour Matrigel HUVEC tube formation assay. A. Brightfield images and B. total network length of HUVEC networks after 12 hours of culture in EGM-2 media, basal (DMEM) media, or conditioned media generated by hASCs cultured in DMEM media (hASCs-CTRL). C. Images and D. total network length of HUVEC networks generated in conditioned media collected from VEGF or DBCO-VEGF treated hASCS or eASCs.

Table S1. Custom NanoString gene panel to assess ASC regenerative potency ^1-3^.

| **Gene Name** | **HUGO Gene** | **Full Name** | **Probe NSID** | **Category** |
| --- | --- | --- | --- | --- |
| ALPL | ALPL | Alkaline Phosphatase | NM_000478.3:2065 | Osteogenic |
| BGLAP | BGLAP | Bone Gamma-Carboxyglutamate Protein | NM_199173.4:44 | Osteogenic |
| BMP2 | BMP2 | Bone Morphogenic Protein 2 | NM_001200.2:1515 | Osteogenic |
| BMP7 | BMP7 | Bone Morphogenic Protein 7 | NM_001719.1:525 | Osteogenic |
| COL1A2 | COL1A2 | Collagen Type I Alpha 2 Chain | NM_000089.3:2635 | Osteogenic |
| FGFR2 | FGFR2 | Fibroblast Growth Factor Receptor 2 | NM_000141.4:2204 | Osteogenic |
| IGF2 | IGF2 | Insulin Like Growth Factor 2 | NM_000612.4:765 | Osteogenic |
| IHH | IHH | Indian Hedgehog Signaling Molecule | NM_002181.2:1693 | Osteogenic |
| MMP9 | MMP9 | Matrix Metallopeptidase 9 | NM_004994.2:1530 | Osteogenic |
| RUNX2 | RUNX2 | RUNX Family Transcription Factor 2 | NM_004348.3:1850 | Osteogenic |
| SEMA3A | SEMA3A | Semaphorin 3A | NM_006080.1:585 | Osteogenic |
| SMAD5 | SMAD5 | SMAD Family Member 5 | NM_005903.5:1044 | Osteogenic |
| SOX9 | SOX9 | SRY-Box Transcription Factor 9 | NM_000346.2:2135 | Osteogenic |
| SP7 | SP7 | Sp7 Transcription Factor (Osterix) | NM_001173467.1:1510 | Osteogenic |
| OPN | SPP1 | Secreted Phosphoprotein 1 (Osteopontin) | NM_000582.2:760 | Osteogenic |
| TNFRSF11B | TNFRSF11B | Osteoprotegerin | NM_002546.2:1075 | Osteogenic |
| WNT16 | WNT16 | Wnt Family Member 16 | NM_057168.1:1621 | Osteogenic |
| WNT5a | WNT5A | Wnt Family Member 5A | NM_003392.3:475 | Osteogenic |
| CCL2 | CCL2 | C-C Motif Chemokine Ligand 2 | NM_002982.3:123 | Immunomodulatory |
| CCL7 | CCL7 | C-C Motif Chemokine Ligand 7 | NM_006273.2:120 | Immunomodulatory |
| MCSF | CSF1 | Colony Stimulating Factor 1 | NM_000757.4:823 | Immunomodulatory |
| IL8 | CXCL8 | Interleukin 8 | NM_000584.2:25 | Immunomodulatory |
| HGF | HGF | Hepatocyte Growth Factor | NM_000601.4:550 | Immunomodulatory |
| IDO1 | IDO1 | Indoleamine 2,3-Dioxygenase 1 | NM_002164.5:369 | Immunomodulatory |
| IL10 | IL10 | Interleukin 10 | NM_000572.2:622 | Immunomodulatory |
| IL1RN | IL1RN | Interleukin 1 Receptor Antagonist | NM_000577.3:480 | Immunomodulatory |
| IL6 | IL6 | Interleukin 6 | NM_000600.3:364 | Immunomodulatory |
| GAL9 | LGALS9 | Galectin 9 | NM_002308.3:359 | Immunomodulatory |
| PTGS2 | PTGS2 | Prostaglandin-Endoperoxide Synthase 2 | NM_000963.1:495 | Immunomodulatory |
| TSG6 | TNFAIP6 | TNF-Stimulated Gene 6 Protein | NM_007115.2:250 | Immunomodulatory |
| TNFRSF11A | TNFRSF11A | TNF Receptor Superfamily Member 11a (RANK) | NM_003839.3:226 | Immunomodulatory |
| TNFSF11 | TNFSF11 | TNF Superfamily Member 11 (RANKL) | NM_003701.2:490 | Immunomodulatory |
| ANGPT1 | ANGPT1 | Angiopoietin 1 | NM_001146.3:2080 | Angiogenic |
| VEGFA | VEGFA | Vascular Endothelial Growth Factor A | NM_001025366.1:1325 | Angiogenic |
| VEGFB | VEGFB | Vascular Endothelial Growth Factor B | NM_003377.3:687 | Angiogenic |
| GAPDH | GAPDH | Glyceraldehyde-3-Phosphate Dehydrogenase | NM_001256799.1:386 | Housekeeping |
| GUSB | GUSB | Glucuronidase Beta | NM_000181.3:1899 | Housekeeping |
| OAZ1 | OAZ1 | Ornithine Decarboxylase Antizyme 1 | NM_004152.2:313 | Housekeeping |
